# High-throughput Phenotyping of Soybean Biomass: Conventional Trait Estimation and Novel Latent Feature Extraction Using UAV Remote Sensing and Deep Learning Models

**DOI:** 10.1101/2024.05.14.594109

**Authors:** Mashiro Okada, Clément Barras, Yusuke Toda, Kosuke Hamazaki, Yoshihiro Ohmori, Yuji Yamasaki, Hirokazu Takahashi, Hideki Takanashi, Mai Tsuda, Masami Yokota Hirai, Hisashi Tsujimoto, Akito Kaga, Mikio Nakazono, Toru Fujiwara, Hiroyoshi Iwata

## Abstract

High throughput phenotyping serves as a framework to reduce chronological costs and accelerate breeding cycles. In this study, we developed models to estimate the phenotypes of biomass-related traits in soybean (*Glycine max*) using unmanned aerial vehicle (UAV) remote sensing and deep learning models. In 2018, a field experiment was conducted using 198 soybean germplasm accessions with known whole genome sequences under two irrigation conditions: drought and control. We used a convolutional neural network (CNN) as a model to estimate the phenotypic values of five conventional biomass-related traits: dry weight, main stem length, numbers of nodes and branches, and plant height. We utilized manually measured phenotypes of conventional traits along with RGB images and digital surface models from UAV remote sensing to train our CNN models. The accuracy of the developed models was assessed through 10-fold cross-validation, which demonstrated their ability to accurately estimate the phenotypes of all conventional traits simultaneously. Deep learning enabled us to extract features that exhibited strong correlations with the output (i.e., phenotypes of the target traits) and accurately estimate the values of the features from the input data. We considered the extracted low-dimensional features as phenotypes in the latent space and attempted to annotate them based on the phenotypes of conventional traits. Furthermore, we validated whether these low-dimensional latent features were genetically controlled by assessing the accuracy of genomic predictions. The results revealed the potential utility of these low-dimensional latent features in actual breeding scenarios.

## 1. Introduction

Measuring and modeling plant growth in response to biotic and abiotic stresses is crucial for developing stress-robust cultivation management systems and breeding stress-tolerant varieties [1, 2]; however, detailed monitoring of plant traits, such as the number of branches (NB), number of nodes (NN), main stem length (SL), and aboveground weight, presents challenges as the plant grows. Conventionally, these traits have been assessed destructively through plant harvesting, which precludes the continuous monitoring of individual plants. Recently, high throughput phenotyping (HTP) has emerged as a promising technique for plant growth measurement, offering nondestructive measurement, data analysis, and modeling through machine learning integration [3]. HTP facilitates the estimation of phenotypic values for target traits by applying machine learning models to data collected via nondestructive methods, including imaging and 3D measurement of plants. For instance, remote sensing using unmanned aerial vehicles (UAVs) has become a prominent approach to data collection in the field [4, 5]. Such UAVs are equipped with various sensors, including RGB cameras [6, 7], multispectral cameras [8], and Light Detection and Ranging (LiDAR) devices [9, 10]. Although RGB cameras are commonly employed because of their ease of use, affordability, and minimal maintenance requirements, multispectral cameras are becoming popular for measuring vegetation indices associated with growth and yield [11].

The aboveground biomass of plants serves as a critical indicator for assessing the nutritional status and yield potential of crops [12–14]. It represents the total amount of photosynthetic products and thus directly correlates with plant growth, yield, and soil nitrogen levels. In recent years, various models have been proposed to estimate the phenotypic values of the aboveground biomass in soybean (*Glycine max*) by leveraging the relationships between UAV remote sensing images and manually collected data. For instance, Maitiniyazi et al. [15] developed a stepwise multiple regression model to estimate soybean dry weight (DW) based on predicted values of plant height (PH) and volume, and Mohsen et al. [16] utilized 35 vegetation indices obtained via UAV remote sensing to predict soybean fresh weight using ensemble learning techniques such as deep neural network and random forest. Similar models have been developed for other crops, such as wheat (*Triticum aestivum*) [17], rice (*Oryza sativa*) [18], and maize (*Zea mays*) [13, 19], which typically require multiple flights or expensive multispectral cameras or LiDAR systems. In contrast, our approach facilitates phenotypic estimation using data from a single UAV equipped with an inexpensive RGB camera.

While conventional machine learning approaches frequently require manual feature engineering to construct models [13, 14], deep learning frameworks automatically perform feature engineering based on conventional data [20]. Leveraging this advantage, deep learning has seen widespread adaptation to the HTP of plants over the past decade [21]. Various models have been proposed to evaluate phenotypes in images to address challenges such as object detection, classification, and regression. Object detection models have been used to count the number of flower buds and fruits for yield prediction [22, 23], and classification models have been employed to detect plant diseases, determine disease severity, and assess vegetable and fruit quality [24–26]. Regression models have also been used to estimate crop maturity and leaf number [27, 28]. Of note, all three model types offer more accurate phenotypic measurements than conventional machine learning models.

Plant phenotyping can be conceptualized as the condensation of higher-dimensional information on plant morphology. When visually assessing plant morphology, its multidimensional information is typically condensed into lower-dimensional predefined features. Similarly, hand measurements using destructive methods condense high-dimensional information on plant morphology into lower dimensions, including measurable features such as PH and NB. HTP, based on images, offers a different approach for condensing high-dimensional information on plant morphology. In this case, high-dimensional information is translated into RGB images or digital surface models (DSMs), from which low-dimensional features are extracted and measured using machine learning models.

Ubbens et al. [3] introduced latent space phenotyping (LSP) as an innovative extension of HTP. This concept involves defining low-dimensional features derived from high-dimensional data, such as plant images captured by sensors and cameras, as phenotypes within a latent space using dimensional reduction methods. These low-dimensional features are then used instead of conventional phenotypes such as agronomic traits [29]. Principal component analysis (PCA) has been utilized extensively for dimension reduction [30–32]. Similar to PCA, LSP is based on the straightforward principle of utilizing low-dimensional features obtained through automatic dimensional reduction methods applied to collected high-dimensional data. This approach offers greater flexibility than the common HTP methods that typically focus on measuring conventionally defined features. Ubbens et al. [3] utilized low-dimensional features extracted by an embedding network, a type of deep learning model, for a genome-wide association study and quantitative trait locus analysis to identify single nucleotide polymorphisms (SNPs) and loci associated with drought resistance. Interestingly, they found an overlap in the detected SNPs and loci between LSP and previous studies based on conventionally defined features [3]. Similarly, Gage et al. [33] employed PCA and autoencoder to extract low-dimensional features from 3D data of maize acquired using LiDAR in field settings and analyzed them as plant phenotypes. They also calculated the broad-sense heritability of each low-dimensional feature and conventionally defined hand-measured features, discovering that the low-dimensional features were subject to stronger genetic control than the hand-measured features.

While these previous studies on LSP [3, 33] demonstrated the potential of low-dimensional features as selection indices, challenges regarding the interpretability of phenotypes in the latent space were noted as a barrier to their utilization in breeding programs. Ubbens et al. [3] and Gage et al. [33] addressed this issue by employing techniques such as saliency mapping [34] and partial least squares regression to establish connections between conventional descriptors or visible traits and the latent space. In Ubbens et al. [3], saliency maps were used to identify the focus areas of the models from the input data, providing an indirect approach to understanding the latent space. By contrast, in Gage et al. [33], partial least squares regression and its prediction accuracy were used, although this approach may not be universally applicable to all scenarios.

In soybean breeding programs, traits related to aboveground biomass, which is crucial for defining the canopy structure, are key selection targets [34]. Previous studies have focused on developing HTP techniques to estimate traits such as DW, canopy coverage rate, canopy volume, and PH [15, 16, 35]. Despite this progress, a gap remains in the HTP techniques available for capturing the phenotype of specific traits that determine canopy structure, such as NN, NB, and the extent of spread. This deficiency has forced breeders and evaluators to rely on manual field measurements, even though many HTP methods are available. To bridge this gap, we employed convolutional neural network (CNN) models [36] to estimate the phenotypes of traits that define the canopy structure. Our approach made use of the simultaneous analysis of ortho-mosaic images and DSMs of the soybean canopy acquired by UAV remote sensing.

Over the past decade, the throughput of phenotyping has been significantly enhanced by the advent of HTP, which has alleviated the burden of labor-intensive and time-consuming tasks. As Ninomiya [20] suggested, HTP not only mitigates the arduous nature of phenotypic data collection but also opens up new avenues for gathering data on traits defined by novel concepts; however, most previous HTP studies focused on measuring traits conventionally defined by evaluators. With the advent of LSP, there is the potential to harness not only the phenotypes of conventional traits measured in breeding programs but also those traits within the latent space that have eluded conventional measurement techniques. Here, we used CNN models to automatically extract low-dimensional features that represent phenotypes within a latent space. To address the challenge of interpretability, we correlated the extracted features with five conventional phenotypic descriptors: DW, SL, NN, NB, and PH. This was achieved by utilizing the weights of the final layer of the CNN model, which provided a link between the conventional descriptors and the latent space. Furthermore, we assessed the genomic prediction accuracy of these low-dimensional traits and compared them with conventionally defined traits to determine whether the extracted features could serve as viable selection indicators in actual breeding programs.

## 2. Materials and Methods

### 2.1 Field experiment

In 2018, we conducted a field trial for evaluating drought resistance of soybean using a National Agriculture and Food Research Organization core collection consisting of 198 accessions (Supplementally Table S1) in a sandy-soil experimental field at the Arid Land Research Center, Tottori University, Tottori, Japan (35°32’ N lat, 134°12’ E lon, 14 m above sea level). In the field, the soil was covered with white mulching sheets (Tyvek 700 AG, Dupont, USA) to avoid rainwater soaking the ground and to control soil moisture. Drought-stress and control treatments were established using a drip irrigation system. We here refer to the areas of the control and drought-stress treatments as the C-area and D-area, respectively. In the C-area, plants were irrigated three times per day (7:00-9:00, 12:00-14:00, and 16:00-17:00; 5 hours in total). In the D-area, irrigation was stopped after thinning. Sowing was performed on July 3^rd^, 2018, and thinning on July 20^th^, 2018. Fertilizer (15, 6, 20, 11, 7 g/m^2^ of N, P, K, Mg, and Ca, respectively) was added before sowing. Each plot contained four plants of one accession. Each accession was used with two replicates in both the C-and the D-area, and 792 plots were established in total. All plants were planted at 20-cm intervals. Details of these plots are shown in Supplementary Figure S1.

### 2.2 UAV remote sensing

We captured images of the field by UAV remote sensing on August 28^th^ before the destructive investigation. A consumer drone (DJI Phantom 4 Advanced, China) was used for UAV remote sensing, using RGB images of the field immediately before destructive phenotyping. For UAV remote sensing, the field was divided into two parts, and flight plans were set up for each part. The details of each flight plan and shooting conditions for the RGB image were as follows—flight altitude: approximately 12 m; interval for taking pictures: 2 s; focus: autofocus; and white balance: automatic. A flight took approximately 15 minutes and produced 500 to 600 RGB images.

### 2.3 Measurement of phenotypic values in biomass-related traits

PH was measured manually in two plants in each plot on August 21^st^, 2018. The DW, SL, NN, and NB of the respective plants were assessed manually through destructive investigation from September 2^nd^ to 5^th^, 2018. The length of the SL and PH were defined as the distance from the cotyledonary node to the terminal of the main stem and from the surface of the ground to the highest point of the soybean canopy, respectively. These five traits were considered targets for estimation and prediction based on deep learning and genomic prediction (GP). The average phenotypic values of two plants in each plot were calculated as the representative phenotypic values of the plot. Z-score normalization was applied to the phenotypic values of all traits, and the normalized scores were used as the phenotypic values of each plot.

### 2.4 Image preprocessing

A DSM and an ortho-mosaic RGB image of the field were obtained from remote sensing images using software Pix4Dmapper (Pix4D, Switzerland) (Figure 1). The DSM and ortho-mosaic RGB images were segmented for each plot using GPS information. Because non-vegetation regions could be noise in the training of deep learning models, vegetation regions in the DSM and RGB images of each plot were extracted as follows: for the RGB image, the vegetation and non-vegetation regions were segmented using a specified threshold. Pixels with green component values (0–255) that were at least 12% larger than the averages of the red and blue components were considered vegetation regions. Dark pixels with green component values less than10 were removed from the vegetation regions, thus producing masked images of vegetation.

Affine transformation was applied to all obtained images to align the position of the canopy and the orientation of the images. Black pixels were added around the vegetation region to ensure the same spatial resolution for the RGB images of all plots. Consequently, images containing only vegetation regions (height: 390 pixels; width: 520 pixels) were obtained (Supplementary Figure S2).

For the DSM of each plot, the vegetation regions were extracted based on the positions of the vegetation areas in the ortho-mosaic images of the corresponding plots (Supplementary Figure S3). Finally, all preprocessed RGB images and DSMs were manually checked, and images with failed processing results were removed. Images without a canopy in the middle were assumed to be unsuitable for further analysis. The final dataset contained 776 RGB images and DSMs. Image analysis and subsequent deep learning model construction were performed using Python (ver. 3.8.10) (https://www.python.org). OpenCV (ver. 4.4.0.40) (http://opencv.org/) was used for image preprocessing as described above.

To increase the amount of training data for deep learning, data augmentation methods were applied to the ortho-mosaic RGB image and DSM data. The size of the training data obtained from the above procedures was 776, which is smaller than that of the usual applications of deep learning. Because the accuracy of deep learning models generally depends strongly on the size of the training dataset [37], we augmented our dataset. Images were flipped upside down, mirrored, and applied both of them using the *flip* function in OpenCV, which increased the size of the dataset by four times its original size.

**Figure 1.**
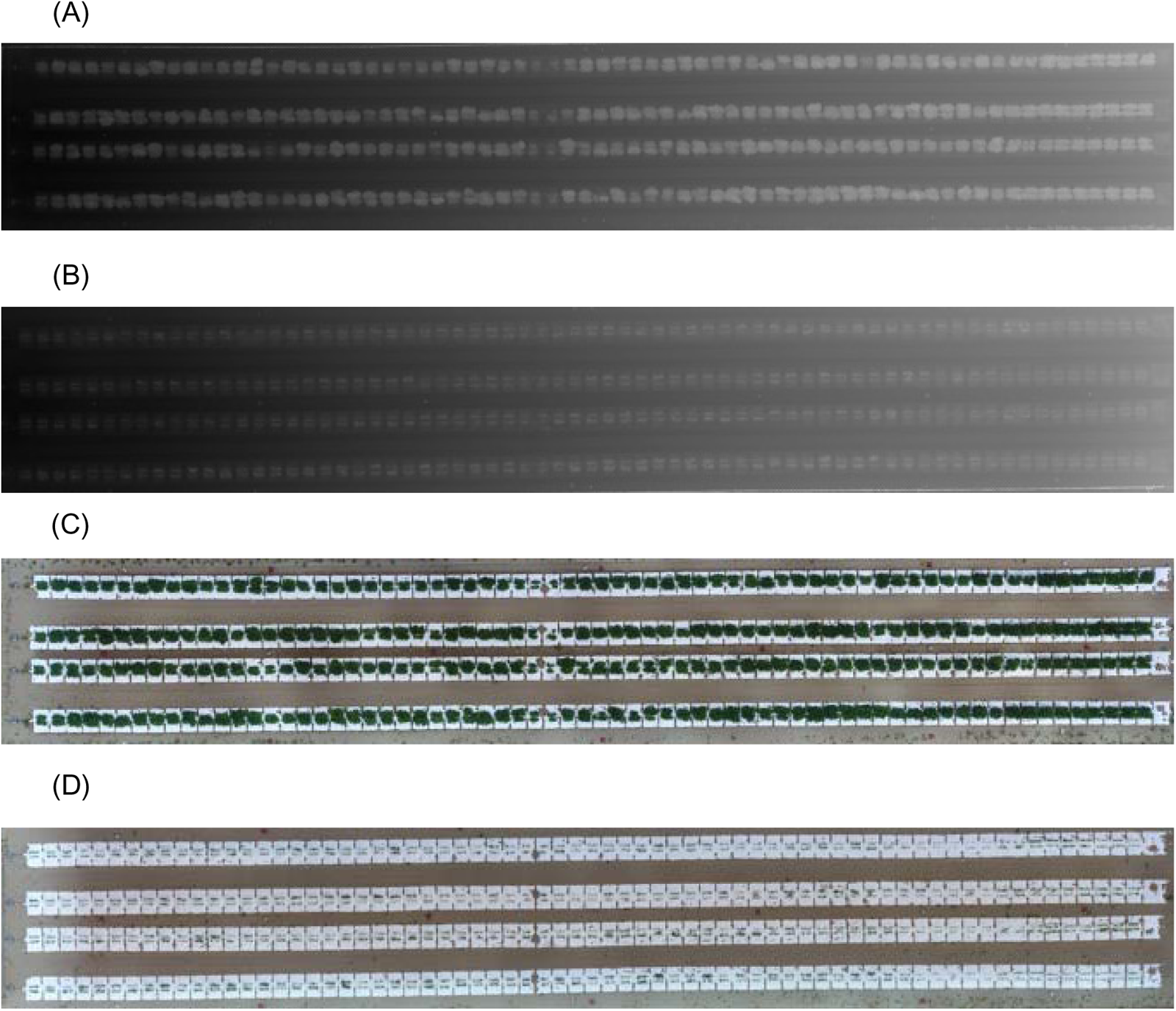
Constructed DSM and ortho-mosaic images by Pix4Dmapper. The color intensity of DSMs illustrates the altitude. The altitude is high as the color is close to white, and it is low as the color is close to black.(A) DSM of the C-area. (B) DSM of D-area. (C) Ortho-mosaic image of the C-area. (D) Ortho-mosaic image of the D-area.

### 2.5 Training of deep learning models

To estimate the phenotypic values of the five target traits (PH, DW, SL, NN, and NB), we constructed deep learning models as CNNs using three different sets of images: (1) RGB images, (2) DSMs, and (3) RGB images and DSMs as inputs. Below, we refer to these three models as RGBNet, DSMNet, and RGB-DSMNet, respectively. The structures of all models are presented in Supplementary Table S2. All models consisted of two main parts: non-linear feature extraction and linear regression. Each model was aimed at simultaneously estimating the phenotypic values of the five target traits. In this process, a 30-dimensional vector was obtained from the non-linear feature extraction part and then used as an input for a linear regression model estimating the five target traits in the regression part. Pytorch (ver. 1.6.0) (https://pytorch.org) was used for model construction.

Below, we introduce mathematical expressions for linear regression. The phenotypic value of *k*th sample of *i*th trait was estimated by the following equation (Equation 1).

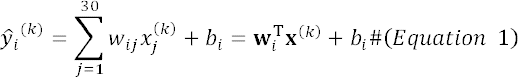

Here, 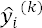 is the estimated phenotypic value of a target trait, 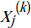 is the *j*th element of the 30-dimensional vector, which are derived from the last layer of the feature extraction part, *w_ij_* is a weight of *x_j_^(k)^* to estimate the phenotypic value of the *i*th trait, *b_i_* is a bias, **w***_i_* is a vector which has *w_ij_* as elements (**w***_i_* = (*w_i_*_1_, …, *w_i_*_30_)^T^), and **x**^(*k*)^ is a vector which has *x_j_^(k)^* as elements 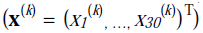. respectively. The values of the 30 elements of *x_j_^(k)^* were scaled such that their means and variances were equal to 0 and 1, respectively. By binding the five equations corresponding to the five target traits, this equation can be written in matrix form:

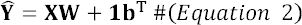

Here, 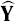 is an *n* x 5 matrix of the estimated phenotypic values 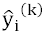 (*n* is the number of samples and 5 is the number of traits) **X** is an *n* × 30 matrix of the extracted features *x_j_^(k)^* , **W** is a 30 × 5 matrix of weights, **1** is a vector of length *n* of which are all 1, and **b** is a vector with *b_i_* as elements (**b** = (*b*_1,_ …, *b*_5_)^T^). To minimize the difference between the observed value **Ŷ** and the estimated value **Y,** the mean square error (MSE) was used as a criterion to determine the optimal values of the regression parameters.

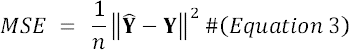

Here, **Y̑** is an *n* × 5 matrix of estimated phenotypic values, **Y** is an *n* × 5 matrix of observed phenotypic values, *n* is the number of samples, and 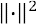 represents the L2 norm. Once the model represented by Equation 1 was constructed, the estimated phenotypic values of the target traits were calculated using the regression parameters optimized using Equation 2, and the features were extracted from the input images using the feature extraction component of the model. The models were trained under the following conditions: number of epochs: 100, initial learning rate: 0.001, batch size: 128, loss function: mean squared error, and optimizer: Adam [38]. The learning rate was adjusted by the *ReduceLROnPlateau* function in *torch.optim.lr_scheduler* module using the following arguments, factor: 0.5, min_lr: 10^−6^, and patience: 5. In each cross-validation repetition, the best model acquired from the training process was saved and used to estimate the phenotypic values of the five target traits. Pytorch was used to train the models and estimate phenotypic values.

The accuracy of all three models (RGBNet, DSMNet, and RGB-DSMNet) for estimating phenotypic values **Y** as **Y̑** was evaluated via 10-fold cross-validation with 20 repetitions. For each cross-validation fold, the dataset was divided into training data (90%) and validation data (10%). In each replicate, the estimation accuracy of the phenotypic values of the five target traits (PH, DW, SL, NN, and NB) was evaluated using the three deep learning models (RGBNet, DSMNet, and RGB-DSMNet) and Pearson’s correlation coefficient. Correlation coefficients were calculated for each trait in each fold of cross-validation, and the averages of these values were used for the final evaluation. We also separately calculated the correlation coefficients for each of the two irrigation conditions (control and drought) to validate the influence of different growth levels on the estimation accuracy of the deep learning models.

### 2.6 Feature extraction from RGB images and DSMs based on RGB-DSMNet

The three deep learning models estimated the phenotypic values of the five target traits in the regression part based on the latent features extracted from the input in the feature extraction part. In other words, each model had a layer consisting of a 30-dimensional vector before the final output layer. We used the values of 30 elements of this vector as latent features. The following analysis was conducted on RGB-DSMNet, which yielded the highest estimation accuracy for most traits, as described in the Results section: Because there is no fixed rule governing the sequence of elements within the deep learning model layer, the interpretation of each element within the 30-dimensional vector varies with each training instance. For instance, the meaning of a specific element within a 30-dimensional vector varies among the iterations of cross-validation. Consequently, considering each element of the vector as an independent variable could disrupt the analysis, intending to regard them as latent features associated with the five target traits.

PCA addresses this issue by determining the axes of the latent feature space, which are spanned by the 30 dimensions of the vector, using a variance maximization criterion. Here, the first 5 or 10 principal components were used to summarize the latent features. The relationship between the 30-dimensional vector and the principal component (PC) scores can be represented by Equation 4:

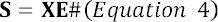

Here, **S** is an *n* × 30 matrix of PC scores for each sample stacked in the row direction, and **E** is a 30 × 30 matrix of the eigenvectors yielding the eigendecomposition of the variance-covariance matrix of **X**. As described above, **X** is an *n* × 30 matrix of the extracted features, and thus, we can calculate the 30 × 30 variance-covariance matrix from **X** because the inverse of **E** is its transpose matrix (**E**^-1^ = **E**^T^):

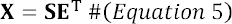

We then visualized the relationship between the PC scores of the 30-dimensional vector and the observed phenotypes of the five target traits. Here, the PC scores of the 30-dimensional vector were treated as low-dimensional representations of the RGB images and DSMs. The weight used to derive phenotypic values (**Y̑**) from PC scores (**S**) can be calculated using the weight of the last layer of the RGB-DSMNet (**W**) and the eigenvectors of **X** (**E**) by assigning Equation 5 to Equation 2:

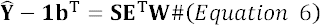

Using this equation, **E**^T^**W** was defined as a new weight matrix to derive phenotypic values scaled based on their means from the PC scores. It was defined as **W’** and used in the visualization process. The sign of the weight exhibited a different pattern in each trial of the 10-fold cross-validation with 20 repetitions. Therefore, the most common sign pattern of the weight matrix was adopted to visualize the relationship between the PC scores of the 30-dimensional vector and the observed phenotypes of the five target traits. The mean value of **W’** in 10-fold cross-validation through 20 repetitions was calculated and used in visualization.

### 2.7 Genomic prediction for biomass-related traits and principal components of latent features

To assess the potential of GP of PCs of latent features, GP models were built for each PC and irrigation level using genomic best linear unbiased predictors (GBLUP) [39]. GP models for the target traits were also built to compare the prediction accuracies of the models. The following model was used for the GP of all traits (DW, SL, NN, NB, PH, and PCs of latent features):

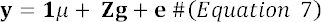

Here, **y** is an *n*-dimensional vector of either the phenotypic values of a target trait or the scores of a PC of the latent features, **1** is a *n*-dimensional vector with all elements 1, μ is the grand mean, **Z** is a *n* × *m* design matrix, **g** is a *m*-dimensional vector of breeding values, **e** is a *n*-dimensional vector of residuals, *n* is the number of samples, and *m* is the number of genotypes. We assumed that **g** and **e** followed the multi-variate normal distribution, 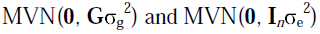, respectively, where **G** is the genomic relation matrix (GRM), 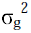, is the additive genetic variance, **I***_n_* is a *n* × *n* identity matrix. and 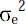 is the residual variance. The GRM was calculated based on SNP genotypes obtained from the whole-genome sequencing data of 198 accessions acquired by Kanegae et al. [40]. Only bi-allelic SNPs were used. SNPs were also filtered by minor allele frequency ≥ 0.025 and missing rate < 0.05. Missing entries in the SNP genotype data were imputed using Beagle 5.1 [41]. As a result, genotypes of 4,776,813 SNPs were obtained and were converted to scores: −1 (homozygous for the reference allele), 1 (homozygous for the alternative allele), or 0 (heterozygous for reference and alternative alleles). The GRM was estimated as **G** = **XX**^T^/c, where **X** is a *n* × *k* SNP genotype score matrix (*n* and *k* are the numbers of genotypes and SNPs, respectively), and *c* is the normalization constant [42], c = 2 Σ*_k’_ p_k’_* (1 – *p_k’_*) where *p_k’_* is the frequency of the allele at marker *k’*.

The accuracy of the genomic prediction model was evaluated using 10-fold cross-validation with 20 repetitions. RAINBOWR package (ver. 0.1.26) [43] was used for the calculations related to GP.

### 2.8 Predicting phenotypic values of traits based on principal component scores of latent features

We performed a further GP to reveal the role of latent features. In this prediction scheme, the phenotypic values of the five target traits were calculated based on the PC scores of the latent features predicted from genomic information. Based on Equation 6, this calculation can be written as

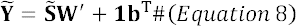

where 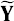 is an *n* × 5 matrix of predicted phenotypic values of five observed traits for *n* samples, and 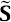 is an *n* × *l* matrix of PC scores predicted by GP. By changing the number of PCs used in prediction (*l*) from 1 to 10, we examined the optimal number of PCs required to predict the genetic variation of each target trait. The number of rows in the weight matrix **W’** is changed corresponding to *l*. The prediction accuracy was evaluated using the same data splitting as in the cross-validation explained in the previous section. To avoid contamination of the test data during the training process, PCA of the latent features was independently performed in every repetition of the cross-validation.

Prediction accuracy was evaluated using Pearson’s correlation coefficient between the predicted and observed phenotypic values for each trait.

## 3. Results

### 3.1 Nondestructive estimation of target traits using deep learning models

All three models showed moderate to high accuracy for the five target traits (Figure 2). The accuracy of PH estimation ranged from 0.881 to 0.946, ranking the highest among all traits evaluated. The estimation accuracy of DW ranged from 0.935 to 0.940. In contrast, the accuracy of the SL, NN, and NB estimations was lower, with correlation coefficients ranging from 0.594 to 0.827. RGBNet demonstrated the lowest estimation accuracy among the three models, except for DW. Conversely, DSMNet provided precise estimates for NB and PH, whereas RGB-DSMNet excelled at estimating DW, SL, and NN.

We also separately calculated the estimation accuracies of the five target traits for each treatment (Figure 3). In all three models (RGBNet, DSMNet, and RGB-DSMNet), the estimation for plants under control conditions was more accurate than that for plants under drought conditions in most cases (except for the estimation results for SL by DSMNet and NB by all three models); however, none of the models could estimate NB under control conditions more accurately than under drought conditions.

**Figure 2.**
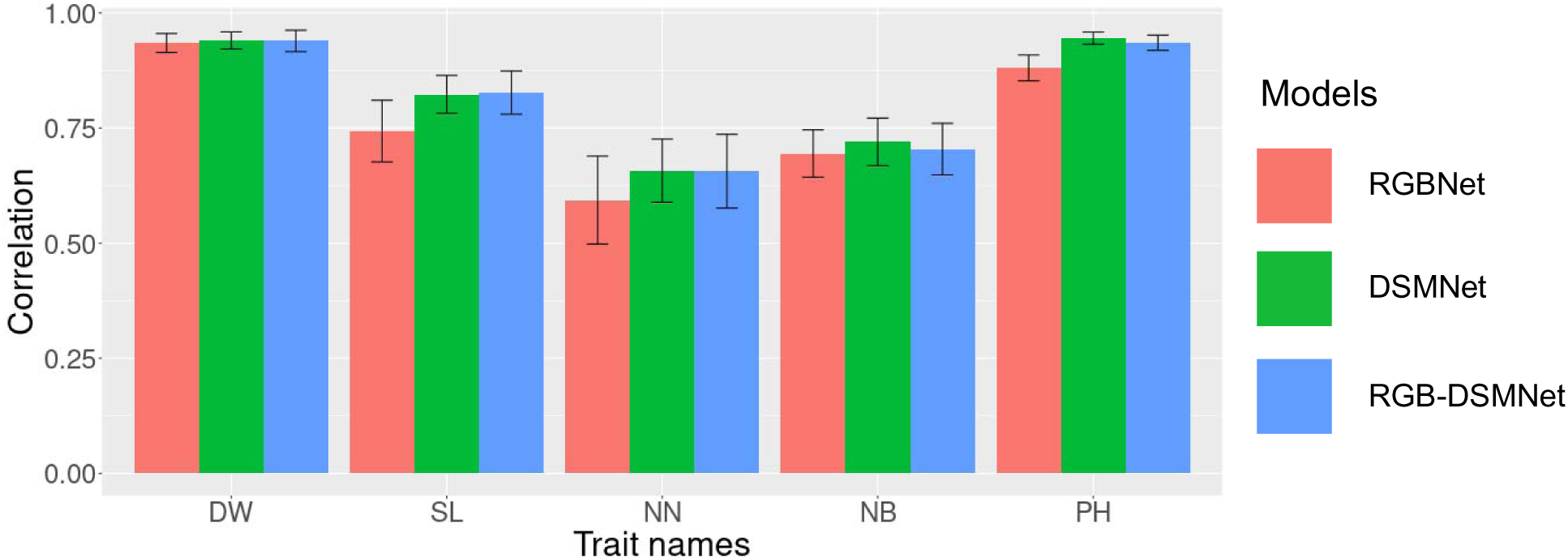
The estimation accuracy of deep learning models. The bars represent the mean value of the correlation coefficient between observed and estimated values obtained through 10-fold cross-validation with 20 repetitions. The error bars indicate the mean standard deviation of the correlation coefficient across the 10-fold cross-validation with 20 repetitions. DW: Dry weight, SL: Length of main stem, NN: Number of nodes, NB: Number of branches, and PH: Plant height.

**Figure 3.**
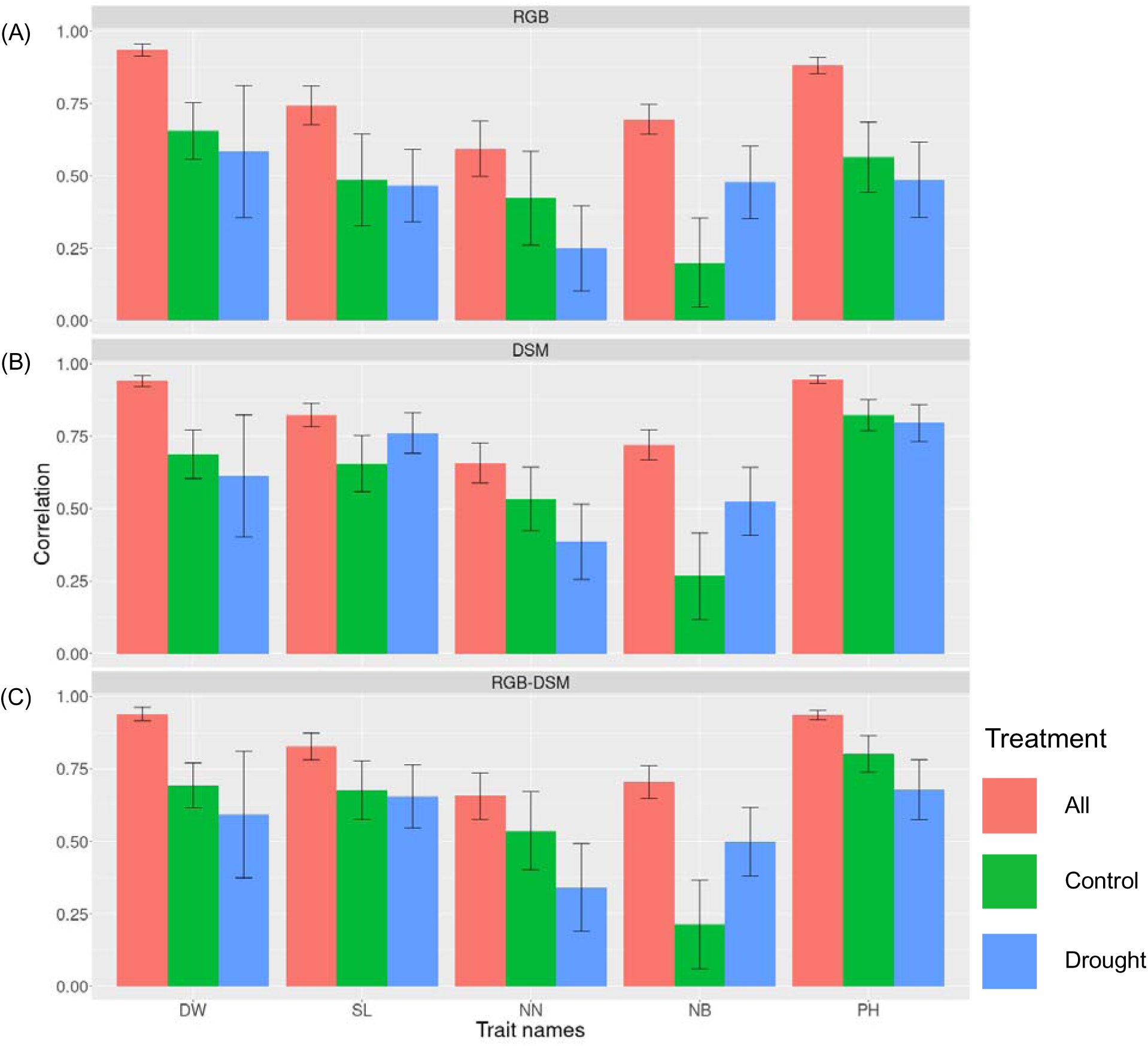
Estimation accuracy of constructed deep learning models under each treatment. The bars represent the mean value of the correlation coefficient between observed and estimated values obtained through 10-fold cross-validation with 20 repetitions. The error bars indicate the mean standard deviation of the correlation coefficient across the 10-fold cross-validation with 20 repetitions. DW: Dry weight, SL: Length of main stem, NN: Number of nodes, NB: Number of branches, and PH: Plant height. (A) Results of RGBNet. (B) Results of DSMNet. (C) Results of RGB-DSMNet

### 3.2 Relationships between latent features and biomass-related traits

The relationships between latent features and biomass-related traits are shown in Figure 4. The relationship between a single trait and the PCs of the latent features varied among the traits. For example, the direction of the effect of the first PC (PC1) on all traits was the same. The direction of effect of PC2 on SL and NN was the same, whereas it was opposite for DW, NB, and PH, compared with the former traits. Likewise, the direction of effect of PC3 on NN, and NB was the same, while it was opposite for DW, SL, and PH, compared with the former traits.

**Figure 4.**
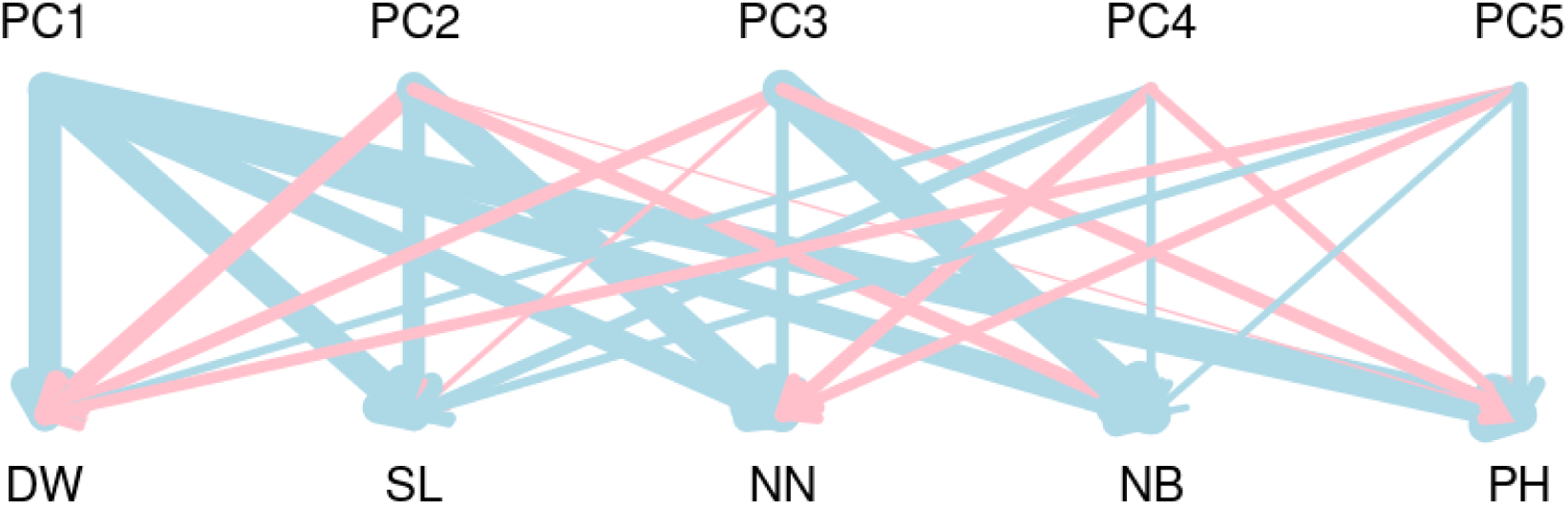
The relationship between principal component of low-dimensional features extracted by RGB-DSMNet. The width of lines represents absolute values of weights between two elements. Blue represents that the values of weights are positive, and pink represents that the values of weights are negative. DW: Dry weight, SL: Length of main stem, NN: Number of nodes, NB: Number of branches, and PH: Plant height.

### 3.3 Genomic prediction in biomass-related traits and principal components of latent features

Genomic prediction models were built for the five biomass-related traits using GBLUP. The prediction models showed higher accuracy under control conditions than under drought conditions for all target traits (Figure 5). The highest accuracy (i.e., correlation coefficient between the predicted and observed values) was 0.880 in NN under control conditions, whereas the lowest was 0.602 in NB under control conditions. The highest accuracy was 0.652 in PH under drought conditions, whereas the lowest was 0.343 in NB under control conditions.

**Figure 5.**
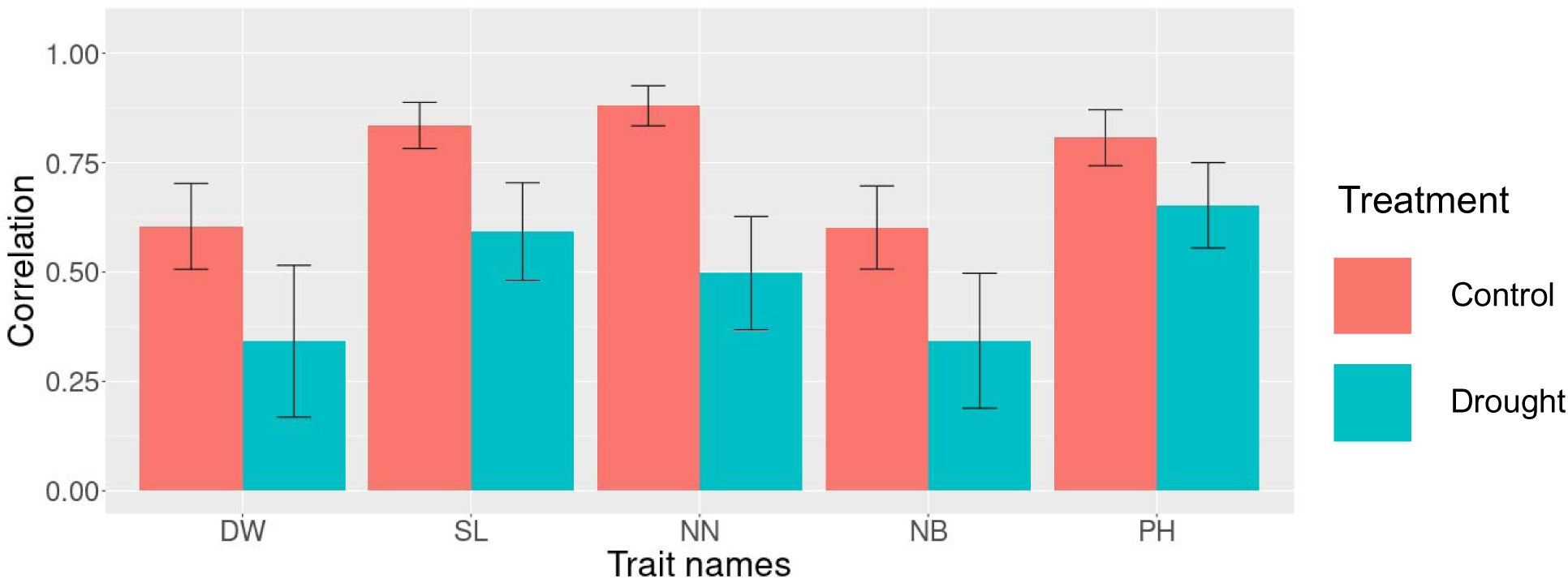
Accuracy of genomic prediction for biomass-related traits. The bars represent the mean value of the correlation coefficient between observed and predicted values obtained through 10-fold cross-validation with 20 repetitions. The error bars indicate the mean standard deviation of the correlation coefficient across the 10-fold cross-validation with 20 repetitions. DW: Dry weight, SL: Length of main stem, NN: Number of nodes, NB: Number of branches, and PH: Plant height.

Genomic prediction models were also built for the PCs of the latent features obtained using RGB-DSMNet (Figure 6). The prediction accuracy was higher under control conditions than under drought conditions for the first three PCs. Under control conditions, the highest accuracy was 0.646 in the first PC, while the lowest was 0.081 in the tenth PC. Under drought conditions, the highest accuracy was 0.281 for the first PC, and the lowest was 0.156 for the tenth PC.

**Figure 6.**
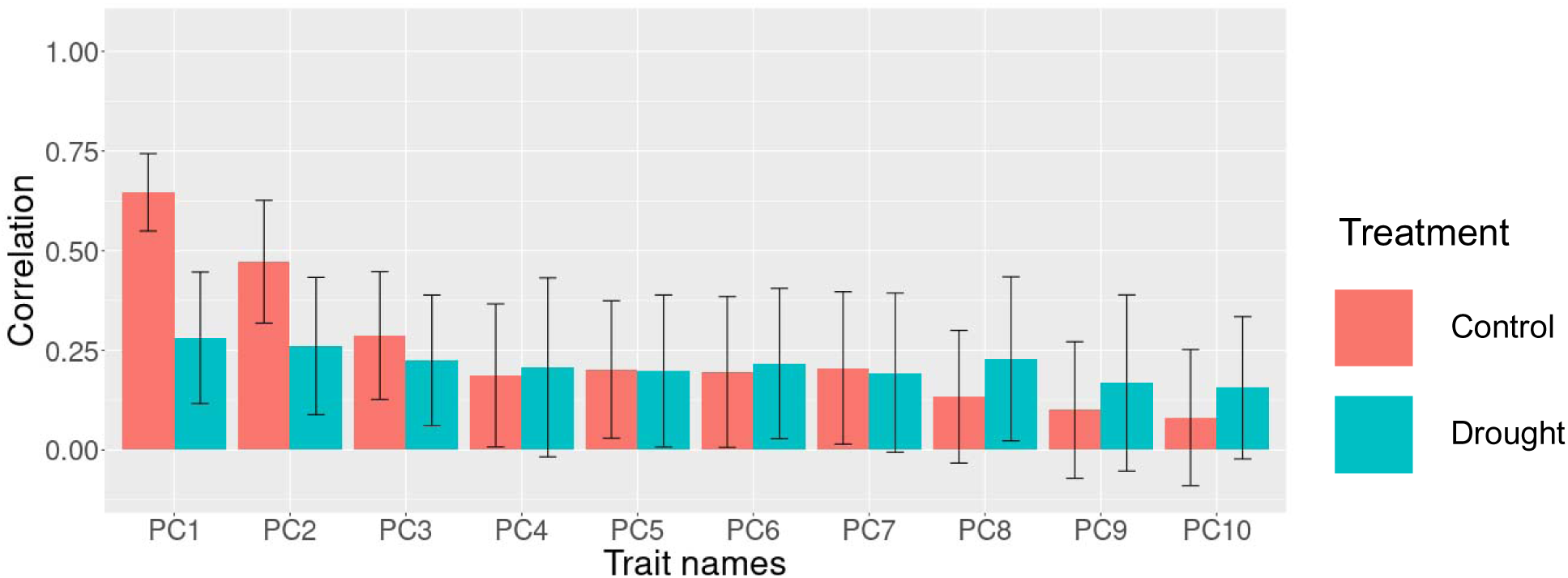
The accuracy of genomic prediction for the principal components of latent features. The bars represent the mean value of the correlation coefficient between observed and predicted values obtained through 10-fold cross-validation with 20 repetitions. The error bars indicate the mean standard deviation of the correlation coefficient across the 10-fold cross-validation with 20 repetitions.

### 3.4 Predicting phenotypic values of biomass-related traits based on predicted principal component scores

The phenotypic values of the five target traits were predicted using the predicted scores of the first ten PCs for genomic prediction. To examine the optimal number of components for prediction, the prediction accuracy was evaluated by increasing the number of PCs (Figure 7). The results showed that predictions based on the first two PCs were as accurate as those based on the first ten PCs for all traits except NB. This suggests that the first two PCs contained essential information for estimating the phenotypic values of the five target traits; however, in NB, an extra PC (i.e., the third PC) seemed necessary. Two different patterns illustrating the relationships between the predicted scores of the PCs and observed phenotypic values of the target traits were confirmed. That is, the first pattern indicates that the first three PCs affected those predictions, while the contributions of the others were almost zero or negative (DW and SL in the control and DW in the drought treatment). By contrast, the second pattern suggested a non-negligible contribution from the fourth and subsequent PCs (NN, NB, and PH in the control and SL, NN, NB, and PH in the drought treatment).

**Figure 7.**
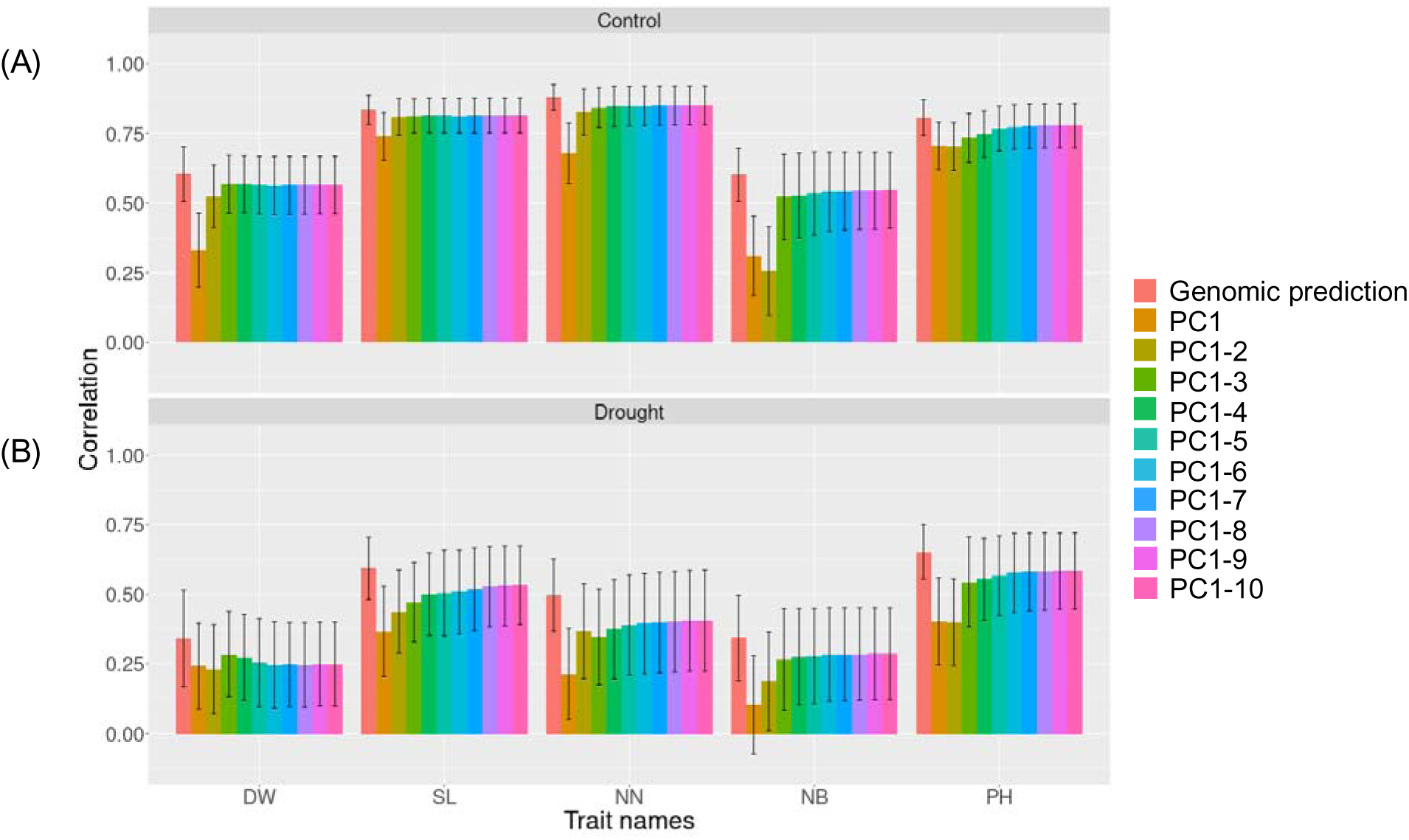
Accuracy of prediction results based on the principal components of latent features predicted by genomic prediction. The bars represent the mean value of the correlation coefficient between observed and predicted values obtained through 10-fold cross-validation with 20 repetitions. The error bars indicate the mean standard deviation of the correlation coefficient across the 10-fold cross-validation with 20 repetitions. ‘Genomic prediction’ indicates that the phenotypic values of all traits were directly predicted by GBLUP. ‘PC1’ indicates that the phenotypic values were estimated based on only principal component score of first principal component predicted by GBLUP, furthermore, ‘PC1-n’ indicates that the phenotypic values were estimated based on principal component score of top n principal components predicted by GBLUP, respectively. DW: Dry weight, SL: Length of main stem, NN: Number of nodes, NS: Number of stems, and PH: Plant height. (A) The results of C-area. (B) The results of D-area.

## 4. Discussion

In the estimation of the phenotypic values of biomass-related traits, DSMNet exhibited higher accuracy than RGBNet across all traits, suggesting that DSMs contain more essential information related to the traits than RGB images. We anticipated that RGB-DSMNet would achieve higher accuracy than the other two models because it utilizes both RGB images and DSMs as inputs; however, while RGB-DSMNet outperformed DSMNet in predicting DW, SL, and NN, as expected, it showed a lower accuracy in predicting PH and NB than DSMNet (Figure 2). This finding suggests that RGB images may be redundant when DSMs are available, as the essential data necessary for the prediction of biomass-related traits within RGB images are already contained in the DSMs.

The RGB-DSMNet can estimate the phenotypic values of DW and SL with high accuracy. Previous studies have developed models to estimate the fresh weight or DW of soybeans because biomass accumulation directly reflects plant growth. Mohsen et al. [16] introduced a deep learning model based on a deep neural network to estimate the fresh weight of the aboveground parts of soybean plants by utilizing 35 vegetation indices derived from data acquired by a hyperspectral camera. Although their model outperformed ours in terms of performance metrics, such as the R-square value, it comes with significant costs for model construction, including a larger number of microplots (approximately 2.5-fold our field experiment) and genotypes (approximately 25% larger than used here), as well as the use of expensive hyperspectral cameras. Their model utilized specific spectral bands related to chloroplasts and plant pigments as input variables. Incorporating data from specific wavelengths into our models could potentially enhance the performance.

Maitiniyazi et al. [15] proposed a stepwise multiple regression model for estimating the DW of the aboveground parts of soybean plants, utilizing PH and volume as architectural data and vegetation indices as spectral data.

A strong correlation between SL and PH may seem intuitive; however, we found that the different models performed best for the destructive estimation of these traits. Specifically, RGB-DSMNet was the top-performing model for SL, whereas DSMNet ranked second.

Conversely, for PH, DSMNet emerged as the best model, followed by RGB-DSMNet as the second-best model. Regarding the NN, the nondestructive prediction was less accurate, with a correlation coefficient between the observed and estimated values below 0.7. This suggests that capturing the relevant information for an NN via remote sensing from above the plant is challenging.

We successfully extracted phenotypic values of biomass-related traits solely from RGB images and DSMs without requiring feature selection, which is typically needed in other machine learning techniques, such as ridge regression, lasso regression, random forest, and support vector machine [44]. This was achieved by leveraging deep learning (CNN). By employing deep learning, we were able to reduce the time and chronological costs associated with the analysis.

The relationship between the PCs of the latent features and biomass-related traits is illustrated in Figure 4, revealing several insights. First, PC1 accounted for all biomass-related traits. Second, the direction of the effect of PC2 on DW and NB was opposite to that on SL and NN. This suggests that plants tend to have a long SL and large NN, but lower NB, shorter PH, and light DW as PC2 increases. In addition, the direction of the effect of PC3 on DW and PH was opposite to that on NN and NB. Hence, it was also inferred that plants had lower DW, shorter SL and PH, but larger NN and NB as PC3 increased. The other PCs exhibited little relationship with the observed traits as their weights were much lower than those of the top three PCs.

The relationship between the latent features and target biomass-related traits was assessed by examining the weights of the output layer of the RGB-DSMNet for PC1, PC2, and PC3; however, this relationship was not apparent for the other PCs because of their significantly smaller weight values. The phenotypic values of the target traits were estimated using the latent features predicted by GP, and their accuracy was evaluated by incrementally increasing the number of components. Consequently, it was suggested that PCs beyond the top three PCs were closely associated with subtle changes that could not be captured by the top three PCs in the target traits, particularly in the case of PH under control conditions, and SL and NB under drought conditions (Figure 7). It is expected that the role of these subtle changes, as illustrated by the PCs beyond the top three, will be revealed and discussed in the field of breeding.

In the study by Ubbens et al. [3], LSP was introduced and utilized to explore quantitative trait loci and SNPs associated with specific traits; however, they did not attempt to annotate the meaning of the phenotypes evaluated as LSP using empirical descriptors such as weight, length, and number. In contrast, Gage et al. [33] defined the minimum requirements for utilizing LSP in an actual breeding program, emphasizing that latent features must be annotated by empirical descriptors and genetically controlled. In our study, the latent features were annotated using the weight of the CNN model, and their genetic control was revealed by the GP results. Our framework acquired the phenotypes of the target traits and annotated the latent features of the plant. Therefore, we strongly anticipate that our technique will introduce a new approach to crop plant breeding.

We investigated the possibility of using the PCs of latent features as new selection indices in breeding fields by comparing the accuracy of genomic prediction for the PCs of latent features with those of conventionally observed biomass-related traits. The accuracy of genomic prediction of the PCs of latent features was generally lower than that of biomass-related traits, with a few exceptions. Notably, under the control conditions, the accuracy of genomic prediction for PC1 was higher than that for DW and NB, suggesting that PC1 was genetically controlled.

As noted by Gage et al. [33], latent features must be related to conventional traits in a visualized manner and genetically controlled for use as selection indices in plant breeding. In our study, PC1 satisfied these conditions, suggesting that PC1 can be used as a new selection index. Furthermore, even if the accuracy of genomic prediction for target traits is low, indicating the low heritability of these traits, the PCs of latent features may contain traits that cannot be conventionally defined. Therefore, depending on the target of the breeding program, it is possible to utilize PCs with low heritability.

The breeding cycle has significantly shortened over the past decade, largely owing to the emergence of genomic selection based on GP, HTP, and the integration of these two techniques [45–47]. Although HTP has made substantial contributions, it frequently requires extensive data storage and time to construct models for phenotyping [20]. By contrast, LSP may offer a faster alternative because it does not always require large datasets or specific models to estimate the phenotype from the data. Consequently, LSP can accelerate the breeding cycle more efficiently than HTP alone. Moreover, the utilization of LSP can reveal features in data that breeders have not previously captured, thereby enhancing the potential of breeding programs [3, 20, 33].

## General

We appreciate the technical staff at the Arid Land Research Center, Tottori University, and Izumi Higashida. We greatly appreciate Dr. Bo Zhang for her advices to improve our manuscript.

## Author contributions

MO: Methodology, software, formal analysis, data curation, writing, original draft preparation, and visualization. YT: Investigation, data curation, and project administration. CB and KH: Methodology and software. HI: Conceptualization, investigation, data curation, writing-review and editing, supervision, project administration, and funding acquisition. AK: Conceptualization, resources, data curation, and funding acquisition. YO, YY, HirT, HidT, MT, MYH, HisT, MN, and TF: Conceptualization, data curation, and funding acquisition. All authors contributed to the manuscript and approved the submitted version.

## Data Availability

The Data supporting the findings of this study are available from the corresponding author, HI, upon reasonable request.

## Supplementary Materials

**Figures S1.**
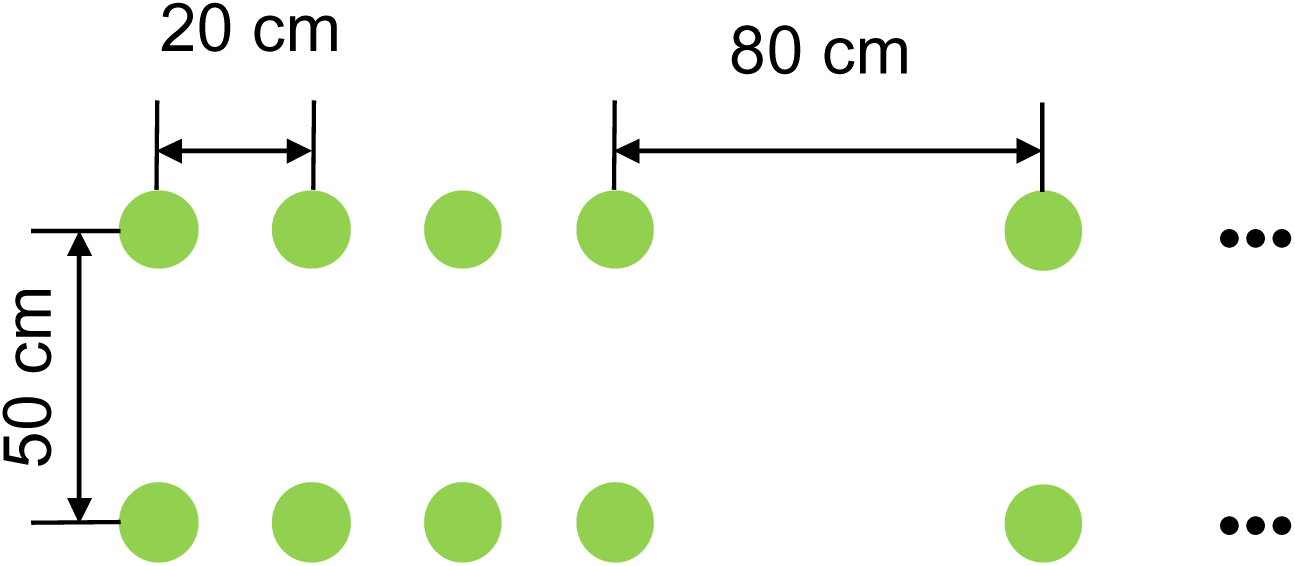
The plot design in each plot

**Figures S2.**
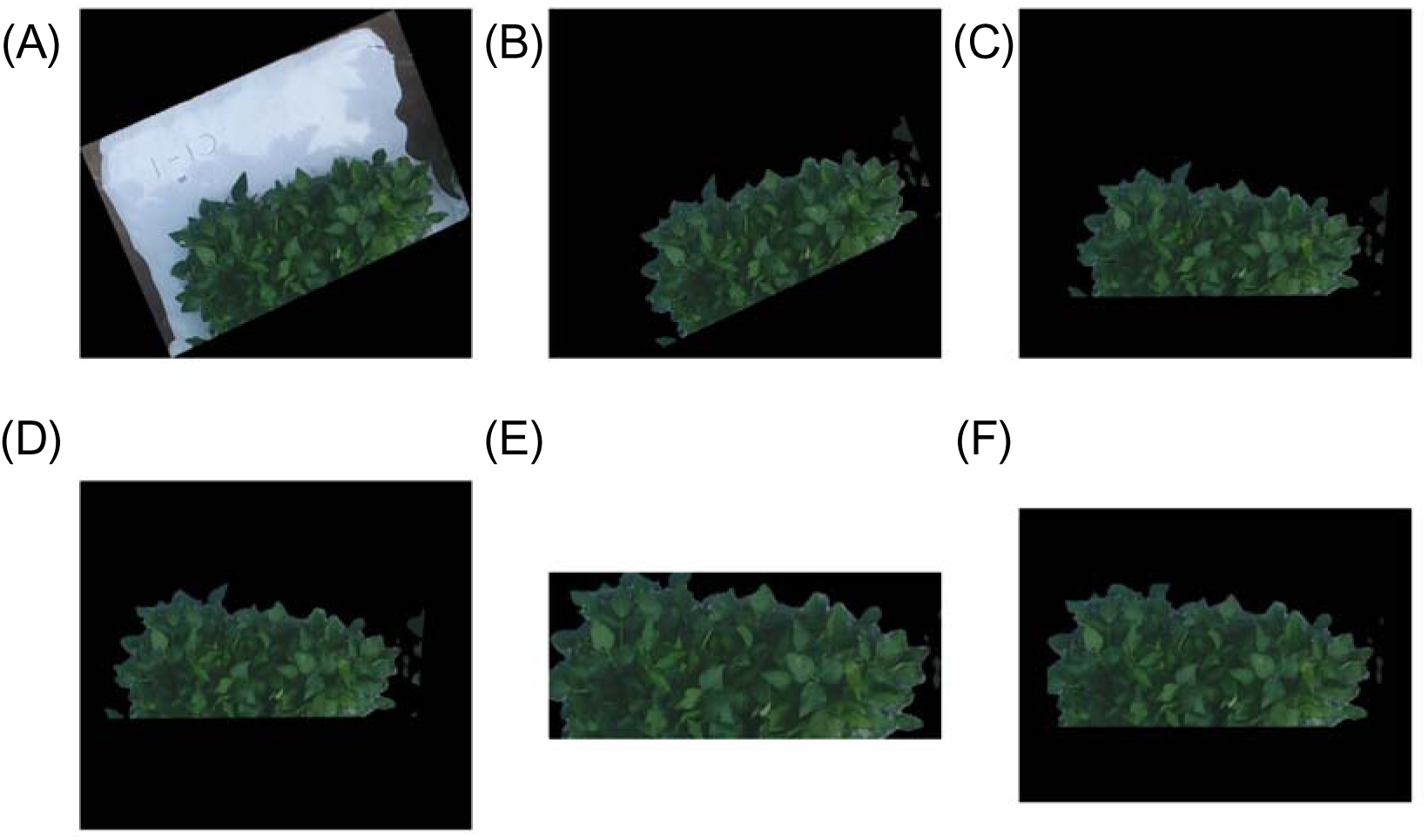
Preprocessing flow of RGB image. (A) Original image of each plot. (B) Masked image generated from original image. (C) Rotated masked image. (D) Centered image. (E) Cropped image. (F) Margin was added for equalizing image size.

**Figures S3.**
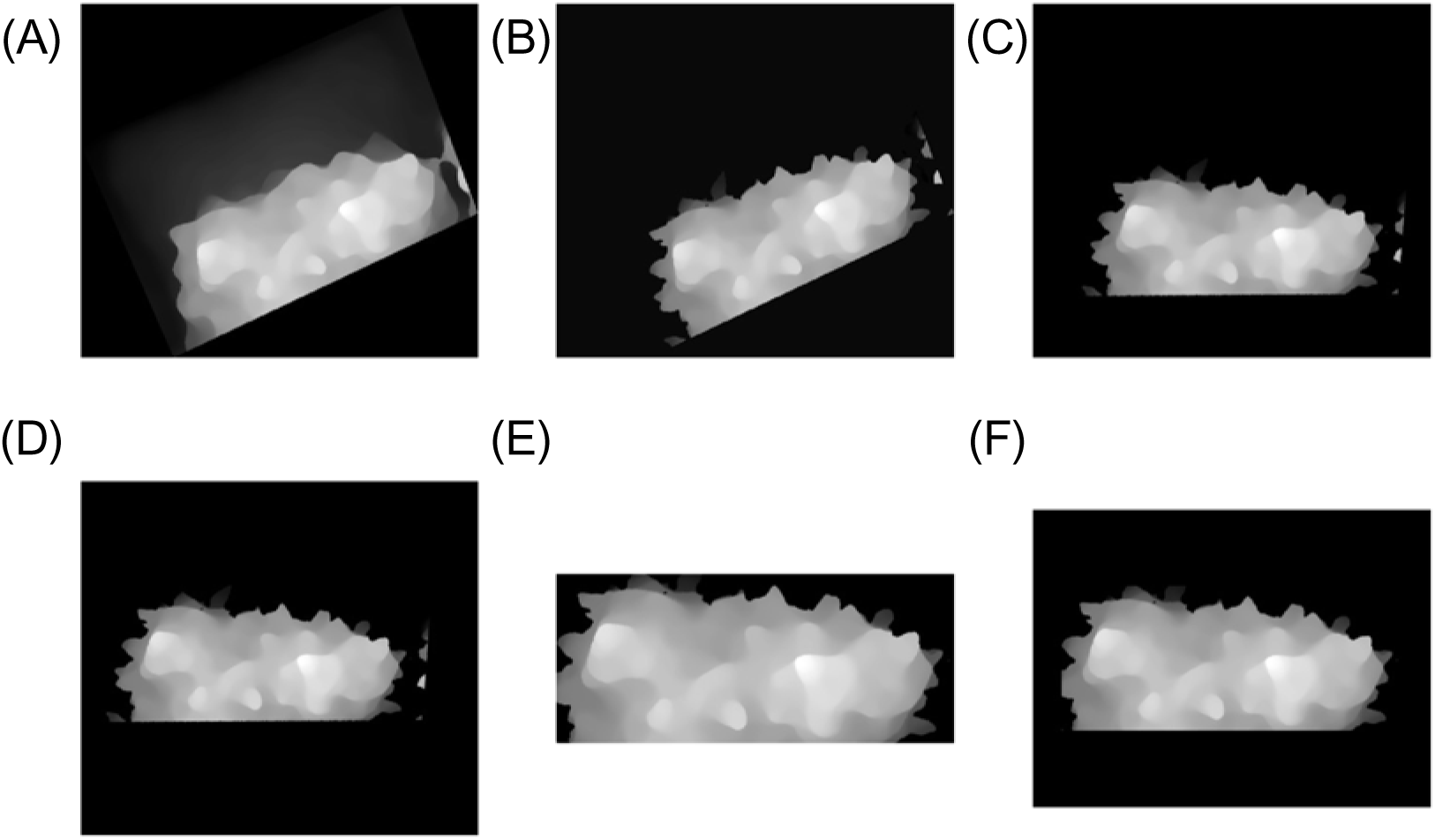
Preprocessing flow of DSM. Matplotlib was used for visualizing the DSM. (A) Original DSM of each plot. (B) Masked DSM generated from original DSM. (c) Rotated masked DSM. (D) Centered DSM. (E) Cropped DSM. (F) Margin was added for equalizing size of DSMs.

