## Supplementary figures and images for "High-throughput Phenotyping of Soybean Biomass: Conventional Trait Estimation and Novel Latent Feature Extraction Using UAV Remote Sensing and Deep Learning Models"

### Supplementary Figure 1

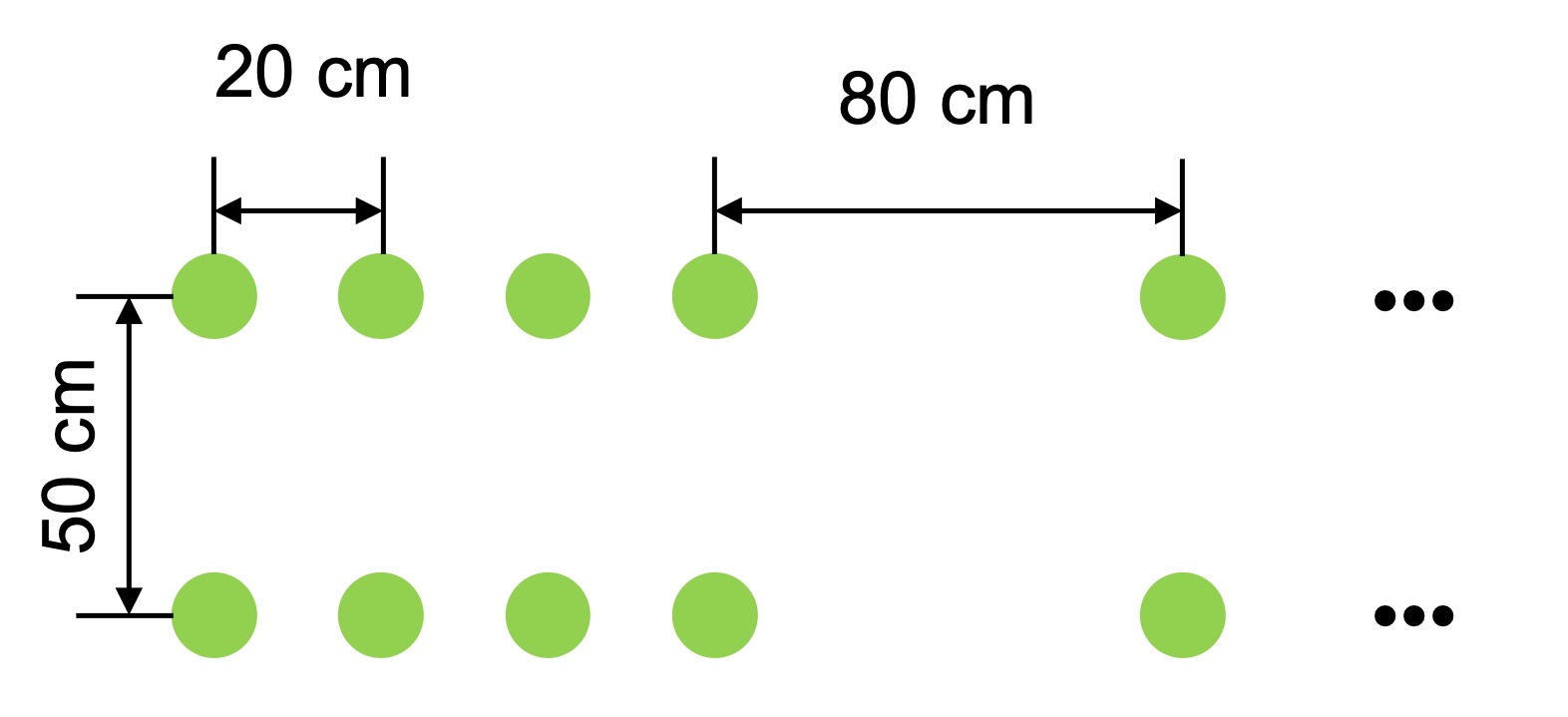

### Supplementary Figure 1

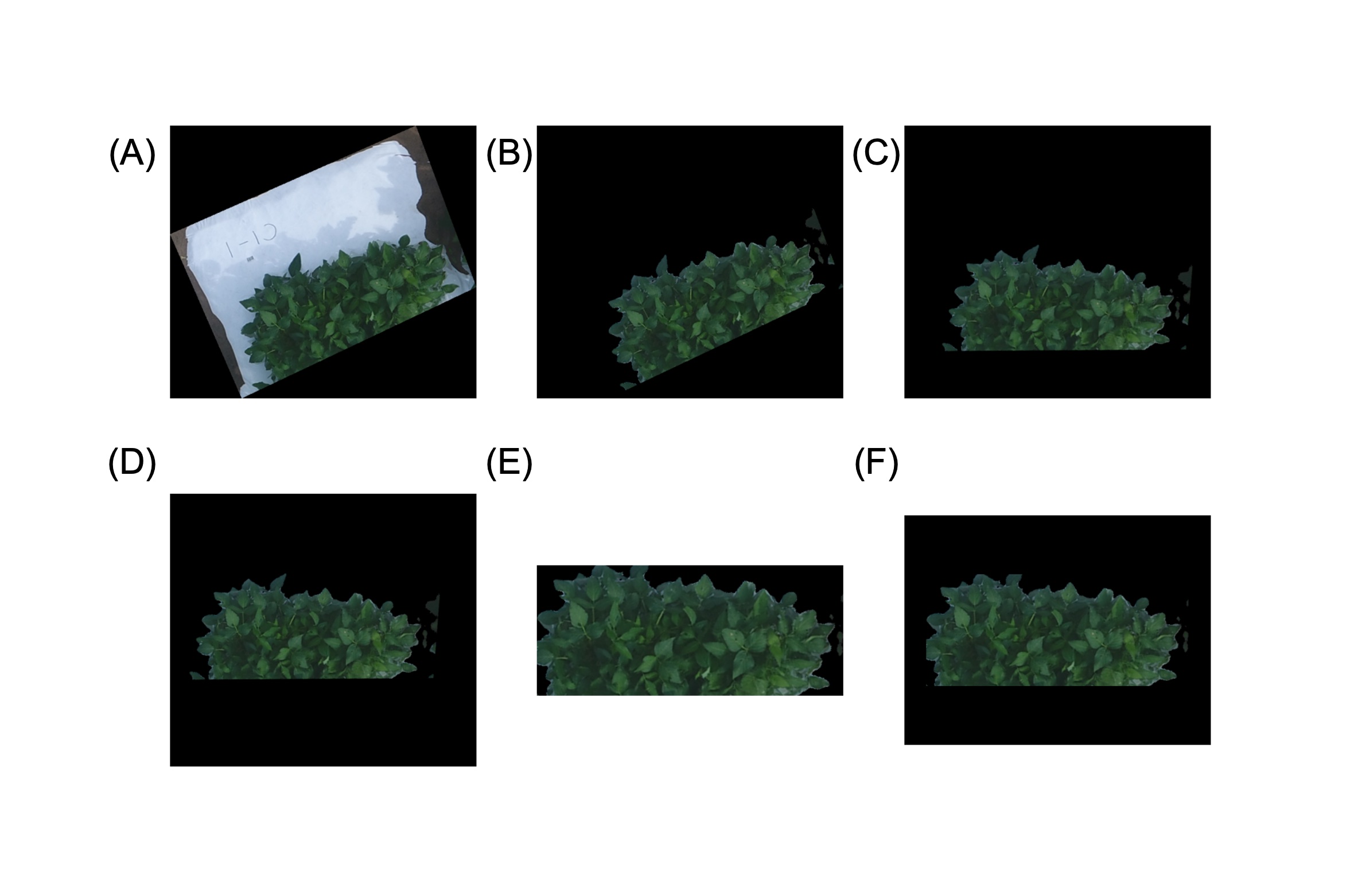

### Supplementary Figure 1

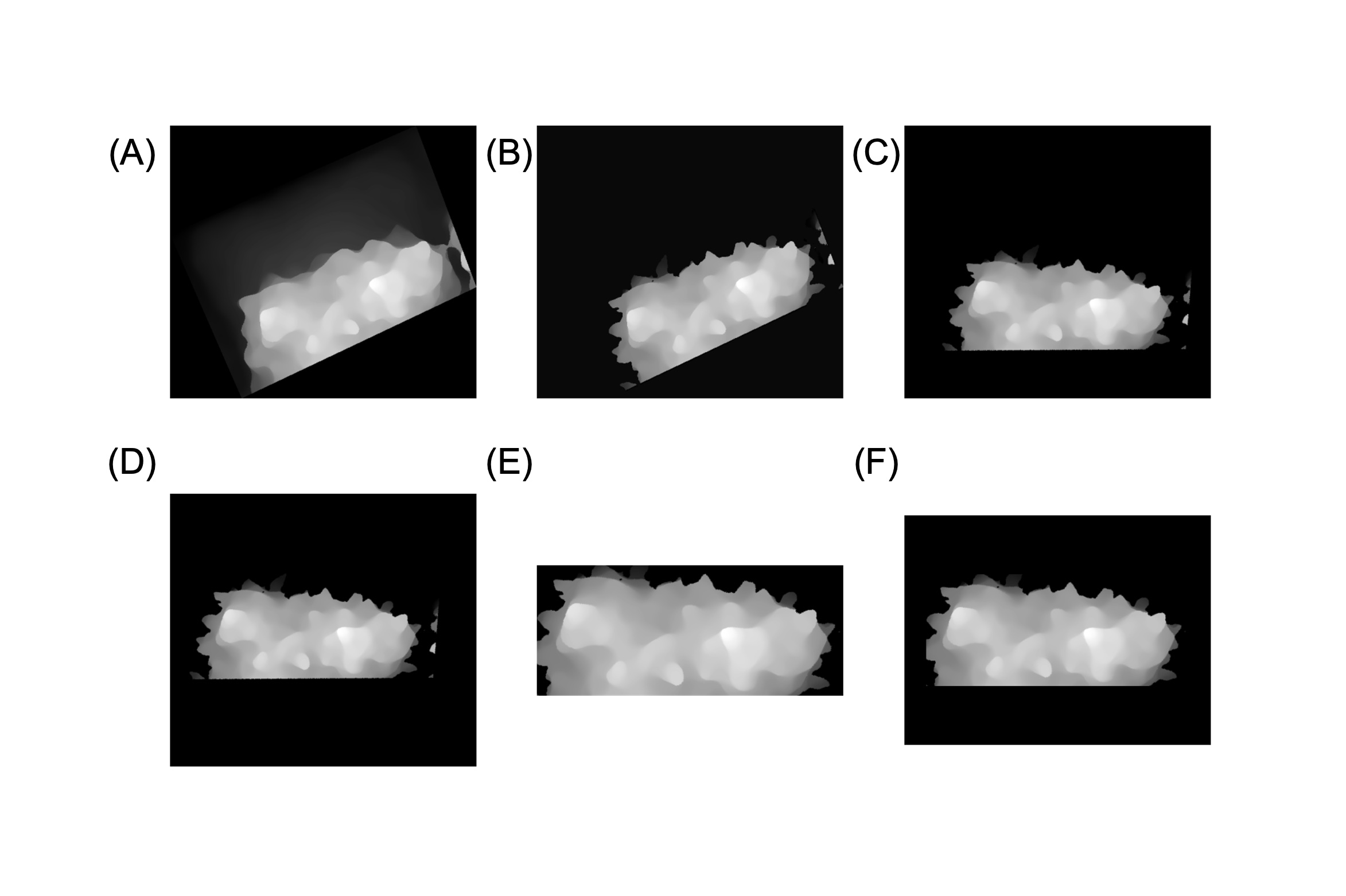
